# Condensate-Driven Triglyceride Depletion Links α-Synuclein to Mitochondrial Dysfunction

**DOI:** 10.1101/2025.10.22.682553

**Authors:** Tao Zhang, María Eugenia Goya, Alejandro Herron-Bedoya, Jorien C. van der Weerd, Dikaia Tsagkari, Suzanne Couzijn, Lale Güngördü, Renée I. Seinstra, M. Rebecca Heiner Fokkema, Ming Chang, Nektarios Tavernarakis, Folkert Kuipers, Ellen A. A. Nollen

## Abstract

Inclusions of α-Synuclein (αSyn) characterize multiple age-related neurodegenerative diseases, including Parkinson’s disease (PD) and Multiple System Atrophy (MSA). While interactions between αSyn and lipids are known to contribute to αSyn pathobiology, the precise cellular mechanisms that link lipids to αSyn toxicity have yet to be elucidated. Through lipidomic profiling of *Caenorhabditis elegans*, we found that αSyn progressively alters lipid metabolism in aging worms. αSyn strongly reduces overall content of triacylglycerols (TAG) and disrupts the structure of lipid droplets (LD). These pathological changes depend on αSyn’s properties to condensate and form inclusions. Apart from lowering TAG levels, αSyn also increases the proportion of long-chain unsaturated fatty acids (LCUFAs). Consequently, genetic inhibition of LCUFA biosynthesis alleviates αSyn-induced loss of *C. elegans* motility. Strikingly, bypassing lipid metabolic defects by supplementing Medium Chain Fatty Acids (MCFAs) restores the αSyn-impaired mitochondrial response and rescues motility. These results link αSyn condensation to impaired TAG metabolism, which reduces mitochondrial function and enhances overall toxicity. Together with the finding that plasma TAGs are lowered in Parkinson patient cohorts, these results suggest that restoring TAG metabolism could alleviate αSyn-induced toxicity in Parkinson’s and other age-related synucleinopathies.

## Introduction

α-Synuclein (αSyn) is an intrinsically disordered protein implicated in the pathogenesis of a number of age-related neurodegenerative diseases, including Parkinson’s Disease (PD), Multiple System Atrophy (MSA), and Dementia with Lewy Bodies (DLB)^1^. Abnormal αSyn conformations and trafficking defects promote formation of proteinaceous inclusions such as Lewy bodies, which disrupt cellular function and eventually lead to disease symptoms^2, 3^. The N-terminal region of αSyn contains seven conserved KTKEGV motif repeats that mediate binding to negatively charged membrane lipids. Recent studies suggest that αSyn-induced toxicity arises from its dual role in disrupting lipid raft architecture and interfering with metabolic pathways^4-6^. Specifically, αSyn penetrates lipid rafts via hydrophobic interactions with the membrane lipids, ultimately forming pores that disrupt the integrity of the membrane^7^. This lipid-based mechanism is consistent with disease-associated mutations in genes such as glucocerebrosidase (GBA)^8^, sterol regulatory element-binding protein 1 (SREBP-1)^9^, and diacylglycerol kinase theta (DGKQ)^10^. However, while *in vitro* interactions between lipids and αSyn have been characterized^6, 11^, downstream impacts on lipid metabolism and resulting consequences on αSyn toxicity remain poorly understood.

Multiple lines of evidence suggest that αSyn toxicity is associated with disruptions in fatty acid saturation and overall lipid composition, both of which are essential for maintaining cellular homeostasis. These disruptions involve changes in monounsaturated fatty acids (MUFAs)^12^, sphingolipids^13^, and phospholipid species^14^. Given the diversity of lipids and their alterations following αSyn condensation, it is clear that disturbed lipid dyshomeostasis is closely associated with αSyn toxicity during development and progression of synucleinopathies. However, whether specific disruptions in lipid metabolism result from the loss of αSyn’s physiological function or from its toxic gain of function remains an unresolved question.

Previously, we developed a transgenic αSyn model in *Caenorhabditis elegans* (*C. elegans)* in which the human αSyn gene, fused to the Yellow Fluorescent Protein (YFP) gene, is expressed in the body-wall muscle cells^15^. This model enables real-time visualization of αSyn condensation, aggregation and age-dependent motility decline, recapitulating aspects of αSyn pathology observed in patients^15-17^. In our previous genetic screen in this model, we identified several modifiers of αSyn toxicity and candidates of interest, in particular genes linked to lipid metabolism^18, 19^. Because central features of lipid metabolism pathways are conserved between humans and *C. elegans*, including shared homologs of human lipid-associated genes^20^, *C. elegans* is a valuable model for identifying the origins of lipid dysregulation and its contribution to age-related diseases^21, 22^.

Here, by performing a lipid metabolism screen in *C. elegans* expressing αSyn-YFP, we identified that αSyn condensation is associated with lower triacylglycerol (TAG) content and an increased proportion of long-chain unsaturated fatty acids (LCUFAs) in the worm. This phenotype is associated with changed properties of αSyn condensates and formation of inclusions that disrupt dynamics of lipid droplet (LD) formation and alter localization of the LD structural protein PLIN-1. Notably, we found that bypassing defects in TAG metabolism by medium-chain fatty acids (MCFAs) feeding rescues mitochondrial function and motility. Taken together, our findings provide insights into cellular mechanisms promoting αSyn toxicity and propose potential therapeutic strategies for alleviating synuclein pathology.

## Results

### αSyn condensation causes changes in lipidome and lowers triacylglycerol levels in *C. elegans*

In previous work, we identified that αSyn initiates self-assembly of amyloid-like condensates in *C. elegans* on day 11 of adulthood (a stage of advanced aging)^16^. Here, we conducted an untargeted lipidomic screen on day 11 αSyn-YFP expressing worms to investigate how αSyn-dependent activity influences lipid metabolism (Fig. 1a). To ensure that we acquired a broad representation of the lipidome, and in particular of moderately and highly apolar lipids, we used the MMC (MeOH/MTBE/CHCl_3_) approach^23^. When we applied unsupervised principal component analysis (PCA), we identified distinct clustering between the lipid profiles of the αSyn-YFP and YFP strains (Fig. 1b), indicating significant lipidome differences. In all, we quantified a total of 18 different lipid classes (Fig. 1c). Consistent with previous related work, we observed an increase in SM levels in αSyn-YFP worms (Fig. 1d)^24^. Intriguingly, our analyses revealed significant lower TAG abundance with a 72% reduction compared to the YFP control (*p < 0.001*) (Fig. 1e).

**Fig. 1.**
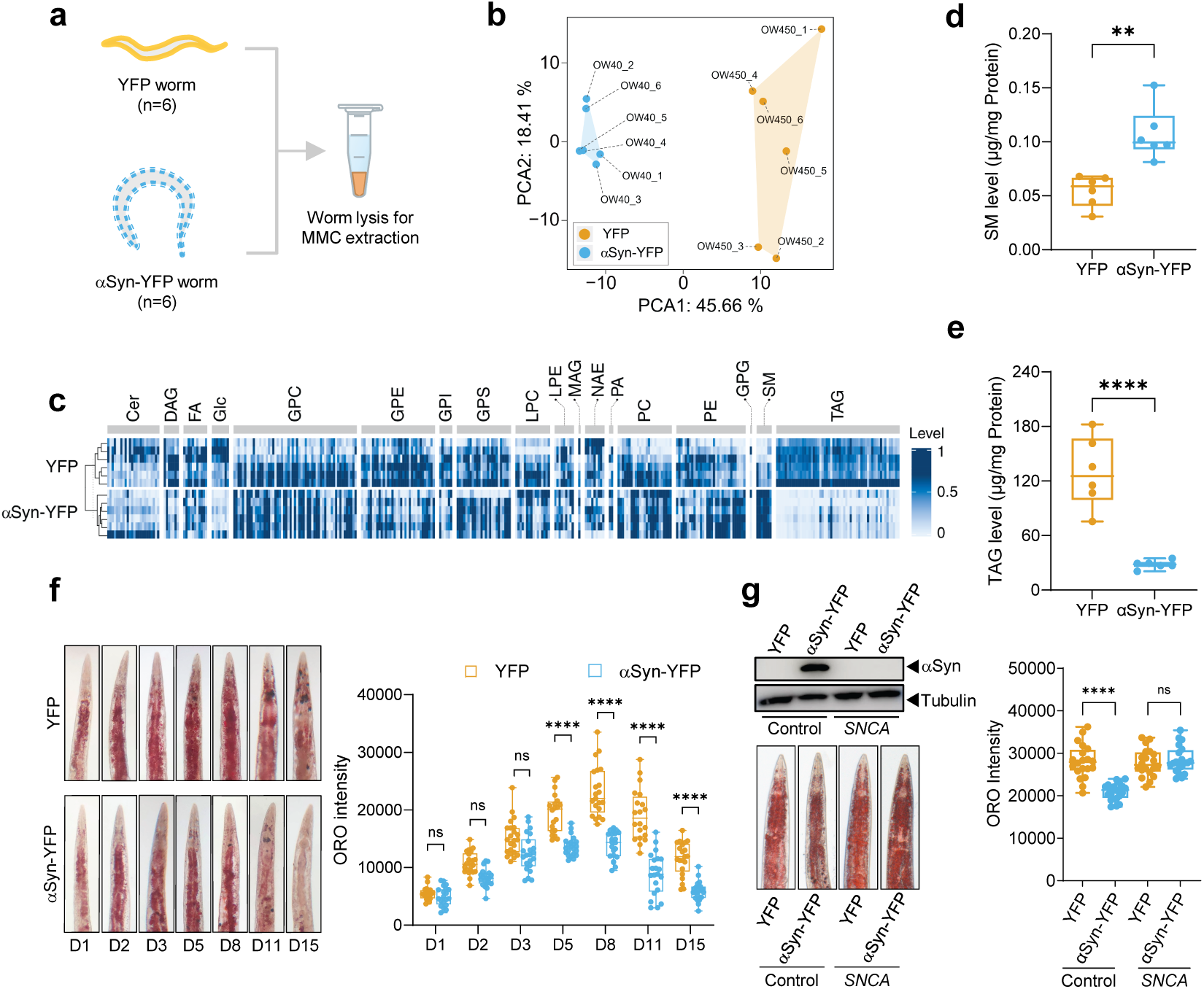
αSyn alters lipid profile and decreases triacylglycerol (TAG) levels in *C. elegans*. ***(a)*** Schematic of sample preparation for untargeted lipidomics. Lipids were extracted by MMC method from whole-worm lysates (YFP and αSyn-YFP strains) at day 11 of adulthood (Each sample contains ∼3000 worms and was collected from six technically replicated plates). ***(b)*** Principal component analysis (PCA) of lipidome data reveals distinct clustering of αSyn-YFP and YFP control worms (n=6). ***(c)*** Heatmap of normalized lipid levels across classes, showing significantly reduced TAG level in αSyn-YFP worms versus YFP controls (n=6). The normalization method employs min-max scaling followed by a log transform to reduce value disparities. ***(d)*** Boxplots of unnormalized sphingomyelin (SM) level in αSyn-YFP and YFP worms (****P < 0.0001, unpaired *t*-test). ***(e)*** Boxplots of unnormalized TAG level in αSyn-YFP and YFP worms (***P < 0.001, unpaired *t*-test). ***(f)*** Oil Red O (ORO) staining confirms decreased TAG level in αSyn-YFP worms from day 5 of adulthood onward compared to YFP controls (n = 20 animals; ****P < 0.0001, unpaired *t*-test). ***(g)*** RNAi knockdown of αSyn (*SNCA*) rescues TAG levels in αSyn-YFP worms at day 5 of adulthood (n = 20 animals; ****P < 0.0001, one-way ANOVA). In *F* and *G*, one representative ORO staining worm is shown in each condition. All experiments were performed in triplicate. TAG: Triacylglycerol, SM: Sphingomyelin, Cer: Ceramide, DG: Diacylglycerol, FA: Fatty Acid, Glc: Glucosylceramide, GPC: Glycerophosphocholine, GPS: Glycerophosphoserine, GPE: Glycerophosphoethanolamine, GPI: Glycosylphosphatidylinositol, LPE: Lysophosphatidylethanolamine, PA: Phosphatidic Acid, MG: Monoacylglycerol, LPC: Lysophosphatidylcholine, NAE: *N*-acylethanolamine, PC: Phosphatidylcholine, PE: Phosphatidylethanolamine, PG: Phosphatidylglycerol.

We next used Oil Red O (ORO) staining to validate the lower TAG level in a time-course analysis (days 1–15). While ORO intensity initially increased and then declined in both groups, staining intensity was markedly lower after day five in αSyn-YFP worms (Fig. 1f), consistent with our lipidomic analysis (Fig. 1c,e). Moreover, as no significant difference in overall ORO intensity was observed at the young adult stage (Fig. 1f), the decreased TAG levels are unlikely to be caused by developmental defects. Taken together, these results align with the age-dependent progression of αSyn toxicity that manifests in later stages of PD and related synucleinopathies.

To assess whether the TAG reduction was due to decreased food intake, we quantified intestinal fluorescence in worms fed mCherry-expressing *E. coli* OP50 (Fig. S1a). As we did not detect any significant difference in fluorescence intensity between αSyn-YFP and YFP control worms (Fig. S1b), we ruled out reduced food consumption as a contributing factor to the lower TAG levels. We also performed RNAi-mediated knockdown of the SNCA gene to investigate whether reduced αSyn expression directly drives the reduction in TAG levels (Fig. 1g). While the intensity of ORO staining in control αSyn RNAi worms was lower than in YFP RNAi worms, depletion of SNCA αSyn restored TAG levels to those of YFP worms (Fig. 1g). Together, these results demonstrate that αSyn causes lipidomic changes and lowers TAG levels in *C. elegans*.

### Reduction in TAG contents mechanistically linked with the formation of αSyn inclusions

Next, we asked whether the reduction in TAG is dependent on αSyn’s properties to form condensates and inclusions. To this end, we assessed TAG levels across wild-type αSyn (WT) and strains expressing two pathogenic αSyn variants (A53T and A30P) associated with familial forms of PD and related synucleinopathies. While αSyn is expressed in all strains, inclusions are formed in worms expressing αSyn and αSyn A53T but not in worms expressing the A30P αSyn variant (Fig. 2a,b). By day 5, TAG levels were significantly lower in αSyn(WT)-YFP and αSyn(A53T)-YFP worms, but unaffected in αSyn(A30P)-YFP worms (Fig. 2c). Pan-neuronal expression of αSyn(WT) or αSyn(A53T) was associated with reduced TAG at day 5 of adulthood (Fig. 2d). These data support that TAG depletion is in part driven by αSyn’s ability to form condensates. In addition, and consistent with its reported stronger neurotoxicity^17^, TAG reduction was more pronounced in neuronal αSyn(A53T)-expressing worms, supporting that the A53T mutation exerts a potent disruption on lipid homeostasis. These results indicate that αSyn-mediated TAG reduction is mechanistically linked to condensation biology, reflecting a fundamental disruption of TAG metabolism.

**Fig. 2.**
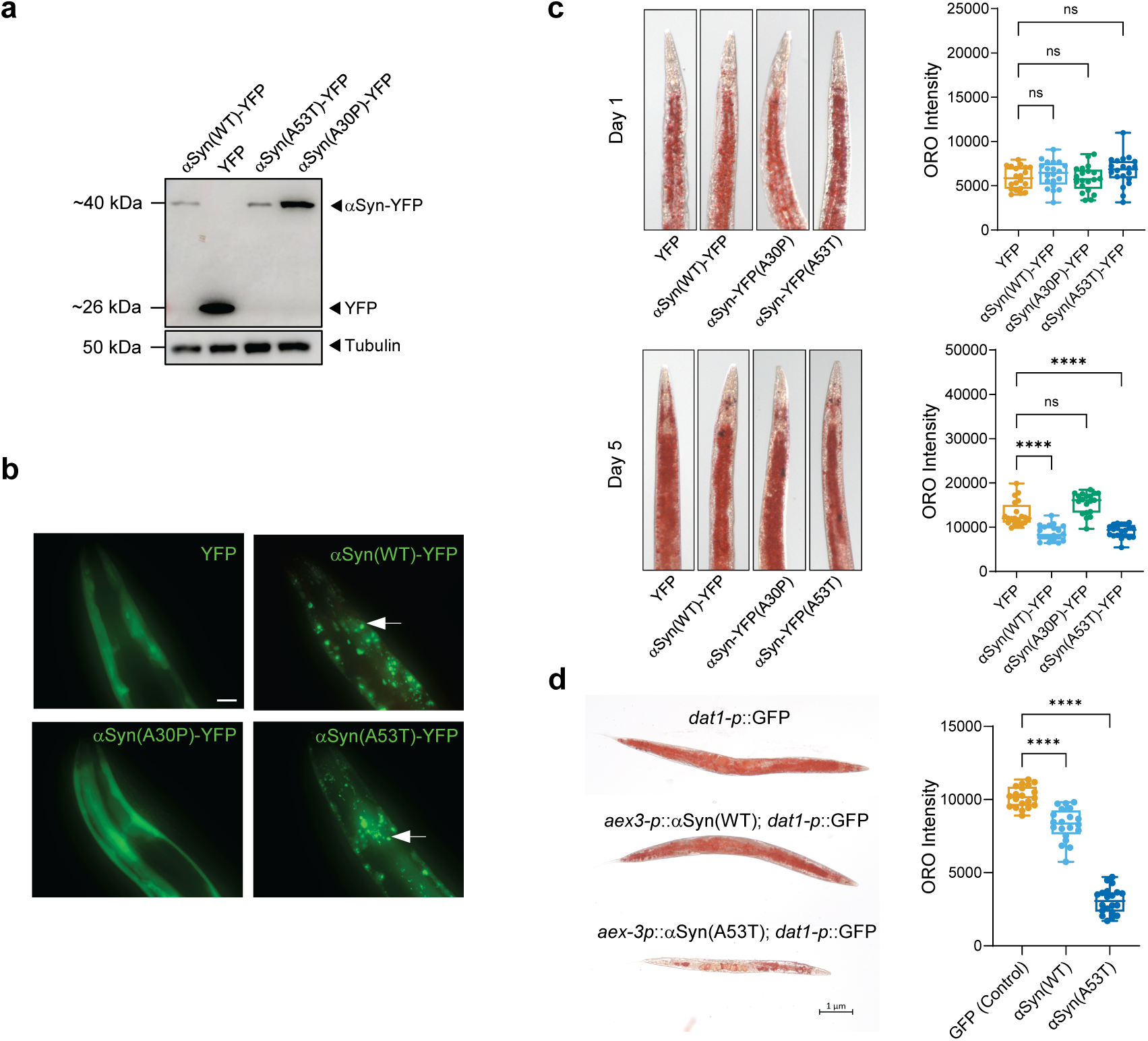
Reduced TAG levels depend on αSyn inclusion-formation. ***(a)*** Western blot analysis of αSyn expression in αSyn(WT)-YFP,YFP, αSyn(A30P)-YFP, and αSyn(A53T)-YFP worms using an anti-GFP antibody. Tubulin was used as control. ***(b)*** Representative fluorescence images showing αSyn inclusions (white arrow) in αSyn(WT)-YFP and αSyn(A53T)-YFP worms, but not in YFP or αSyn(A30P)-YFP worms. Scale bar: 20 µm. ***(c*)** ORO intensity reveals significantly lower TAG levels in inclusion-forming αSyn(WT)-YFP and αSyn(A53T)-YFP worms compared to YFP and αSyn(A30P)-YFP controls at day 5 of adulthood, but no significant difference was observed at day 1 (n = 20 animals per group; ****P < 0.0001, one-way ANOVA). ***(d)*** Pan-neuronal expression of αSyn(WT) or αSyn(A53T) (driven by *dat-1* promoter) reduces TAG levels relative to *dat-1p*::GFP control at day 5 of adulthood (n = 20 animals; ****P < 0.0001, one-way ANOVA). In *B*, *C* and *D*, one representative image is shown in each condition. All experiments were performed in triplicate.

### αSyn reduces lipid droplet abundance and disrupts PLIN-1 localization in muscle

ORO staining reveals overall TAG levels in the worms, and we therefore next questioned whether these are similarly reduced in body-wall muscle cells. TAG is primarily stored in lipid droplets (LDs), which can be visualized using the fluorescent fatty acid analogue Bodipy 558/568 C12. This probe is preferentially incorporated into TAG-rich LDs, allowing detection of LDs^25^. By feeding worms with Bodipy 558/568 C12, we confirmed LD labelling in muscle cells through co-localization with the LD-resident protein DGAT-2, tagged with GFP and expressed under the muscle-specific promoter (myo-3) (Fig. S2a). In αSyn-expressing worms, we identified a striking reduction in muscle LDs at both day 1 and day 5 of adulthood (Fig. 3a,b), suggesting that αSyn-mediated TAG reduction occurs earlier at its primary site of expression.

**Fig. 3.**
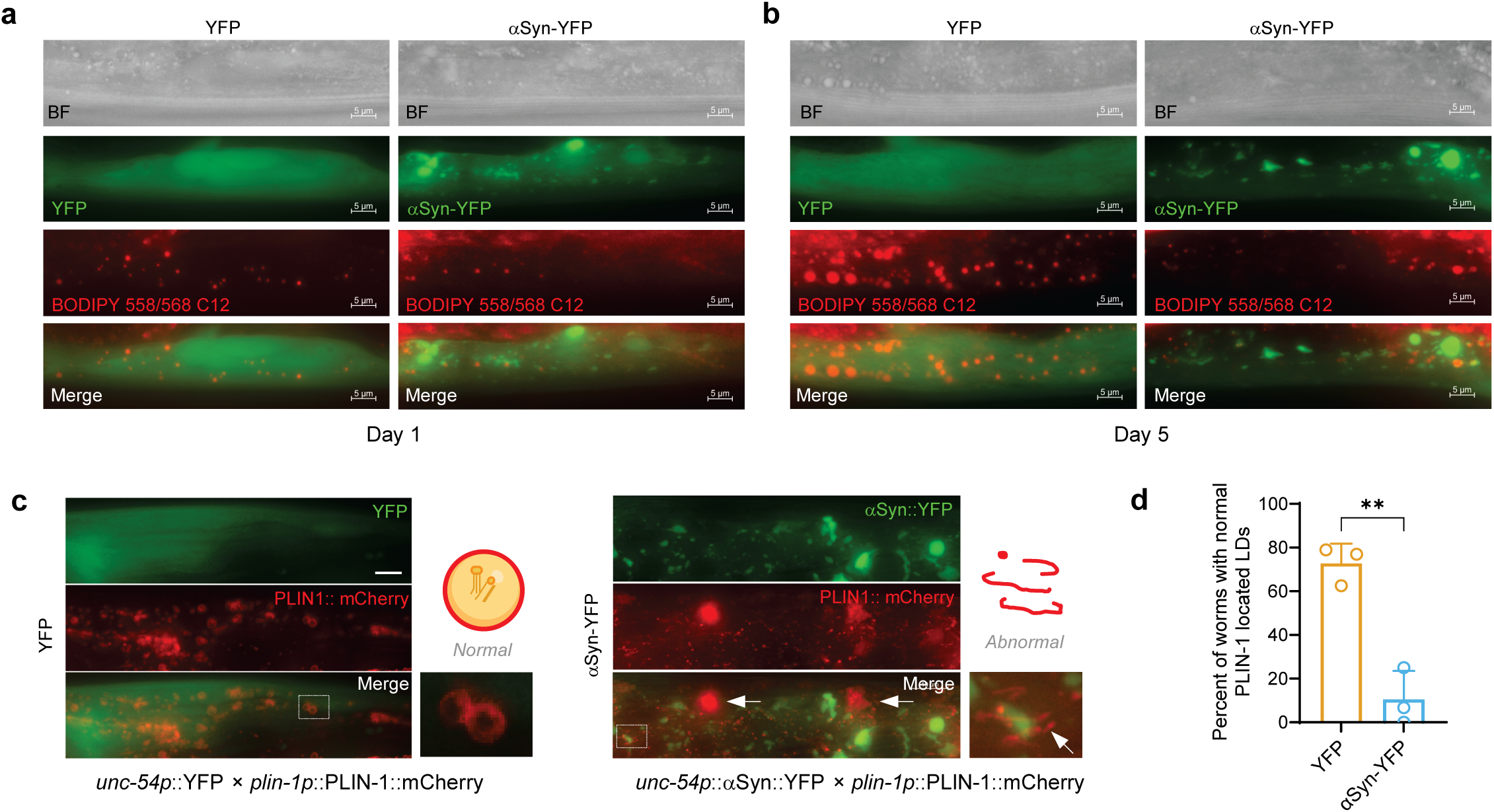
αSyn reduces lipid droplet (LD) abundance and disrupts PLIN-1 localization. ***(a)*** αSyn significantly reduces LD abundance in muscles compared to YFP controls at earlier day 1 of adulthood stage. Scale bar: 5 µm. ***(b)*** αSyn significantly reduces LD abundance in muscles compared to YFP controls at day 5 of adulthood. Scale bar: 5 µm. ***(c)*** Fluorescence reveals αSyn::YFP (green) induces abnormal PLIN-1 distributions including frilly morphologies (white arrow) at day 5 of adulthood. Insets show magnified views of disrupted PLIN-1 localization (white-dotted box regions) and the simplified diagram of disrupted PLIN-1 structure in αSyn-YFP worms. Scale bar: 10 μm. ***(d)*** αSyn inclusion formation decreases the percentage of worms with normal PLIN-1-coated LDs in muscle at day 5 of adulthood (n = 15 animals per group; percentages derived from each biological replicate; **P < 0.01, unpaired *t*-test). All experiments were performed in triplicate.

We next examined whether αSyn condensation affects LD-associated proteins. We focused specifically on PLIN-1, a structural LD-coating protein that regulates lipolysis by controlling fatty acid release from LD-stored TAG for energy supply^26, 27^. We first confirmed expression of PLIN-1::mCherry in muscle by demonstrating its colocalization with DGAT-2::GFP under the muscle-specific promoter (Fig. S2b). Next, we examined the changes in PLIN-1 co-localization. In αSyn-YFP worms, we confirmed that PLIN-1 was severely mislocalized at day 5 of adulthood, with a lower percentage of worms displaying normal PLIN-1-coated LDs compared to YFP controls (Fig. 3c,d). αSyn induces abnormal PLIN-1 distributions as shown in the magnified insets and the simplified diagram of PLIN-1 structure. The abnormal distribution was also more prominent in condensation-prone αSyn(A53T)-YFP worms relative to non-condensating αSyn(A30P)-YFP worms (Fig. S3a,b). Given these results, we hypothesize that a condensate-driven mechanism disrupts proper PLIN-1 localization and LD integrity. Supporting this hypothesis, LDs are smaller in αSyn-YFP worms and with clumpy structures at day 11 of adulthood (Fig. S3c). Notably, physical contact or interaction between αSyn inclusions and LDs is minimal at day 1 of adulthood (Fig. S3d) with no apparent co-localization by day 5 of adulthood (Fig. 3c and Fig. S3a).

These data suggest that TAG depletion and PLIN-1 mislocalization are not solely mediated by direct physical interactions. Instead, these disruptions are more likely to reflect a broader lipid metabolic dysfunction induced by αSyn.

### αSyn-induced reductions in TAG levels and impaired motility are indirectly linked

We also detected significantly decreased motility at day 5 of adulthood in αSyn-YFP worms compared to YFP controls, a phenotype that was rescued by SNCA RNAi (Fig. S4a). To explore the underlying mechanisms, we leveraged our established automated wide-field-of-view tracking platform to perform a targeted RNAi screen for TAG metabolism genes (Fig. 4a)^28^. A schematic representation of the TAG metabolic pathways is shown in Fig. S5. Tryptophan 2,3-dioxygenase (*tdo-2*) RNAi, which was previously shown to enhance motility^29^, served as the positive control, while empty vector (EV) functioned as the negative control. Among all tested genes, we found that knockdown of the stearoyl-CoA desaturases *fat-6* and *fat-7* significantly rescued the impaired motility in αSyn-YFP worms (Fig. 4b). These enzymes play critical roles in polyunsaturated fatty acid (PUFA) biosynthesis and downstream synthesis of other fatty acids and complex lipids. We also observed more modest improvement with knockdown of genes critical for long-chain unsaturated fatty acids (LCUFAs) synthesis, such as *elo-2,* a fatty acid elongase enzyme, and the nuclear hormone receptor *nhr-80*, a known transcriptional regulator of *fat-6* and *fat-7* desaturases^30^. Interestingly, restoration of total TAG content by *atgl-1* RNAi failed to rescue motility (Fig. 4b,c). Overall, these results indicate that the impaired mobility induced by αSyn is independent of the total levels of TAGs but may depend on their properties, such as their length and saturation levels.

**Fig. 4.**
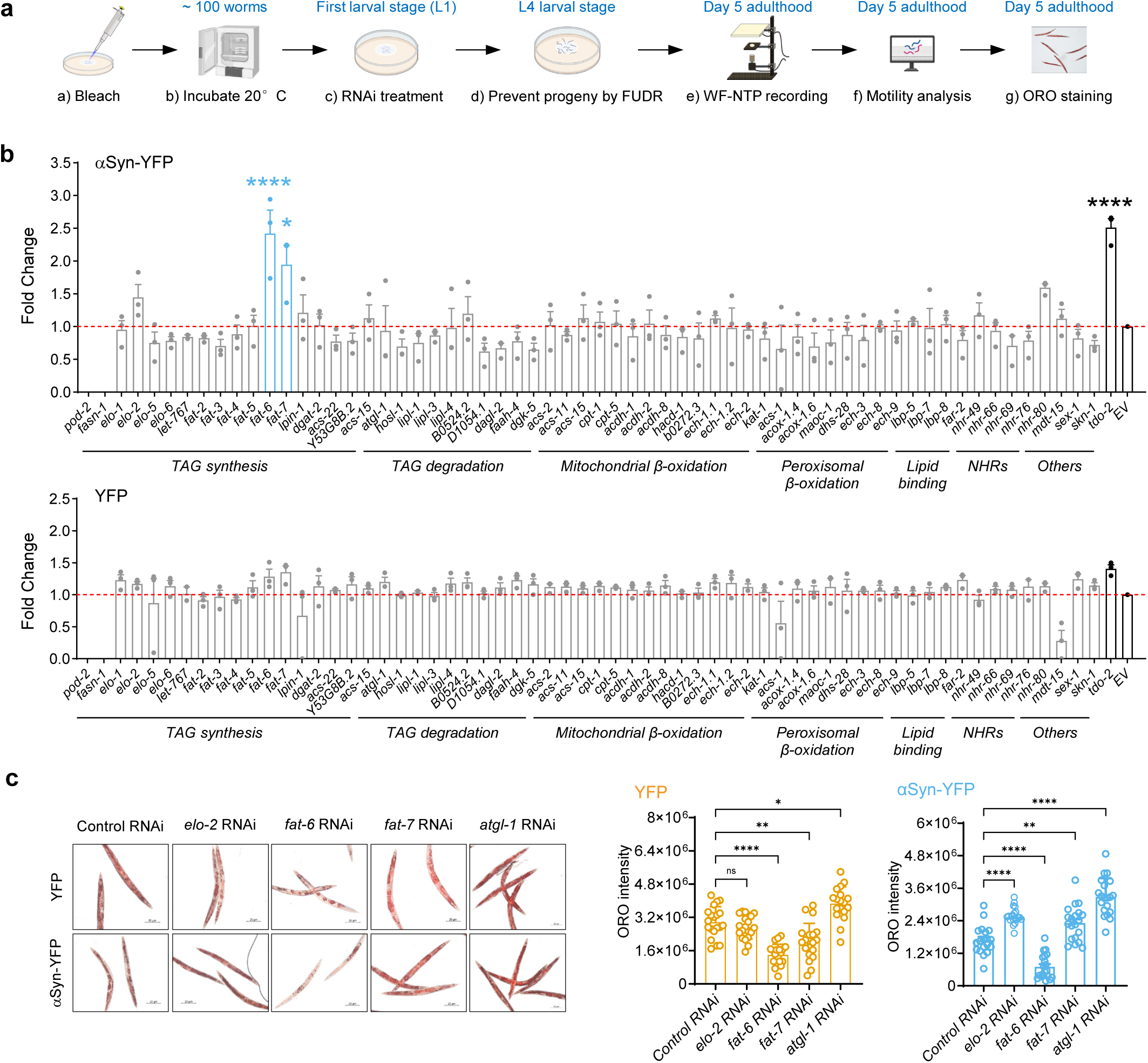
Inhibition of long-chain unsaturated lipid synthesis reduces αSyn toxicity. ***(a)*** Experimental design for RNAi screening. Synchronized L1-stage YFP and αSyn-YFP worms were cultured on RNAi plates at 20°C. FUDR was added at L4 stage to prevent progeny production. Motility screenings were performed in day 5 of adulthood. ***(b)*** RNAi-mediated knockdown of *fat-6* and *fat-7* (blue) significantly reduces αSyn toxicity in day 5 adult αSyn-YFP worms (P < 0.05) but not in YFP control worms. Motility was quantified as body bends per 30 seconds. Empty vector (EV) serves as a negative control and *tdo-2* RNAi as a positive control. ***(c)*** ORO staining shows *fat-6* and *fat-7* RNAi reduce TAG levels in YFP controls at day 5 adulthood, while *fat-7* RNAi and *elo-2* RNAi restore TAG level in αSyn-YFP worms. *atgl-1* RNAi increases TAG in both strains (n=15∼20 animals; *P < 0.05, ****P < 0.0001, one-way ANOVA). One representative image is shown in each condition. Data represent fold changes from biological replicates (*P < 0.05, ****P < 0.0001, one-way ANOVA). All experiments were performed in triplicate.

In contrast, *fat-6* knockdown resulted in significant reduction of TAG levels in both YFP and αSyn-YFP worms. Conversely, *fat-7* RNAi led to restored TAG levels in αSyn-YFP worms (Fig. 4c). The modest improvement in motility observed in YFP worms following *fat-6* RNAi (Fig. 4b), along with the significant reduction in both TAG and αSyn expression levels (Fig. 4c and Fig. S4b), suggest that *fat-6* mediated rescue mechanisms are likely systemic rather than associated with specific suppression of αSyn toxicity. This result is in line with the smaller size of worms following *fat-6* RNAi (Fig. 4c). Thus, *fat-7* may serve as a crucial regulator of αSyn toxicity, influencing both motility and TAG storage. Consistent with this proposed mechanism, *fat-7* overexpression significantly impaired motility. However, a contribution of elevated levels of αSyn in *fat-7* overexpressing worms cannot be excluded (Fig. S4c, d). Together, these findings suggest that *fat-7* likely modulates lipid desaturation to regulate αSyn-induced toxicity.

### Suppressing biosynthesis of long-chain unsaturated fatty acids mitigates αSyn toxicity

Elongation and desaturation of medium-chain saturated fatty acids (MCFAs) into LCUFAs is mediated by *elo-2*, *fat-6* and *fat-7* (Fig. 5a). In our lipidomic analysis, we detected a higher proportion of LCUFAs associated with both TAG and phosphatidylethanolamine (PE), an abundant phospholipid in mammalian cells, in αSyn-YFP worms (Fig. 5b,c and Fig. S6a,b). These data confirm that αSyn toxicity is not solely due to the reduction in total TAG levels, but is also influenced by the specific composition of its fatty acyl chains. When we feed worms with oleic acid (OA), a downstream product of *fat-6* and *fat-7*^20^, the protective effects of *fat-6* or *fat-7* RNAi were abolished (Fig. 5d). Taken together, these results indicate that accumulation of LCUFAs contributes to αSyn-induced pathology and that limiting LCUFAs biosynthesis might mitigate αSyn toxicity.

**Fig. 5.**
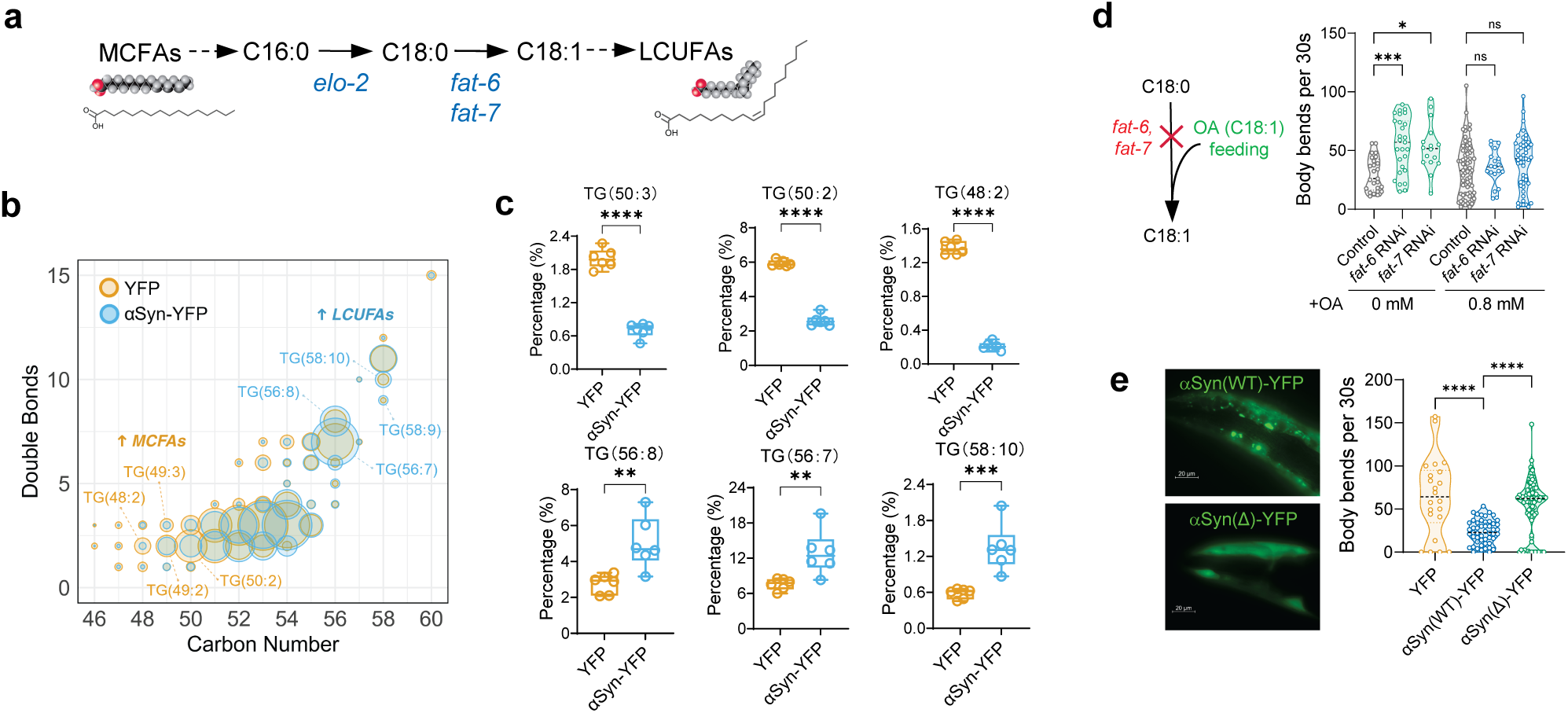
Suppression of long-chain unsaturated fatty acids (LCUFAs) biosynthesis mitigates αSyn toxicity. ***(a)*** Metabolic pathway schematic depicting *fat-6*, *fat-7*, and *elo-2* roles in converting medium-chain saturated fatty acids (MCFAs) to LCUFAs. **(*b)*** Lipidomic analysis reveals αSyn-YFP worms accumulate more LCUFA-containing TAGs (blue) versus YFP controls (MCFAs-enriched, yellow). **(*c)*** Percentage of six predominant TAG molecular species shows significant higher unsaturation in αSyn-YFP worms compared to YFP controls (n=6, **P<0.01, ***P<0.001, ****P<0.0001). ***(d)*** Oleic acid (OA, C18:1) as long-chain unsaturated fatty acid supplementation abolishes the protective effects of *fat-6* and *fat-7* RNAi, failing to rescue motility in αSyn-YFP worms (n=50 animals; *P<0.05, **P<0.01, one-way ANOVA). ***(e)*** Deletion of αSyn membrane-binding domain (PVH198 strain) forms no inclusions and rescues motility defects of αSyn-YFP worms (n=50∼100 animals, ****P < 0.0001, one-way ANOVA). All experiments were performed in triplicate.

It’s widely accepted that LCUFAs physically interact with αSyn in a way that modulates its toxic properties. When we examined the motility of the αSyn(Δ) strain, a transgenic αSyn strain carrying a specific deletion of its membrane-binding domain^31^, we noted significantly higher motility compared to the YFP control (Fig. 5e). These data support that the membrane or fatty acid interactions are a crucial determinant of αSyn-induced toxicity, suggesting that αSyn requires membrane binding to dirsupt lipid metabolism, which impairs LD integrity and reduces motility of the worms.

### Bypassing LCUFAs biosynthesis via MCFAs feeding rescues αSyn toxicity by restoring mitochondrial function

Given that the LD structural protein PLIN-1 is disrupted which serves as a key regulator of LD hydrolysis for mitochondrial energy production, we propose a mechanistic model in which αSyn inclusions disrupt PLIN-1 localization and the mitochondria fragmentation, thereby reducing TAG availability and compromising mitochondrial energy production, ultimately leading to impaired motility (Fig. 6a). Using our established platform for assessment of basal mitochondrial respiration^32^, we found that the mitochondrial respiratory capacity is diminished in αSyn-YFP worms (Fig. 6b), particularly in basal respiratory capacity (Fig. 6c).Additionally,we observed a fragmented pattern in mitochondrial morphology in αSyn-YFP worms, as visualized by TOMM-20::RFP (Fig. 6d). As shown in the magnified inset, we also observed a strong colocalization pattern between αSyn and mitochondria. Taken together, these data show that αSyn accumulates and forms inclusions in the immediate vicinity of mitochondria, contributing to membrane disruption and lower mitochondrial respiration.

**Fig. 6.**
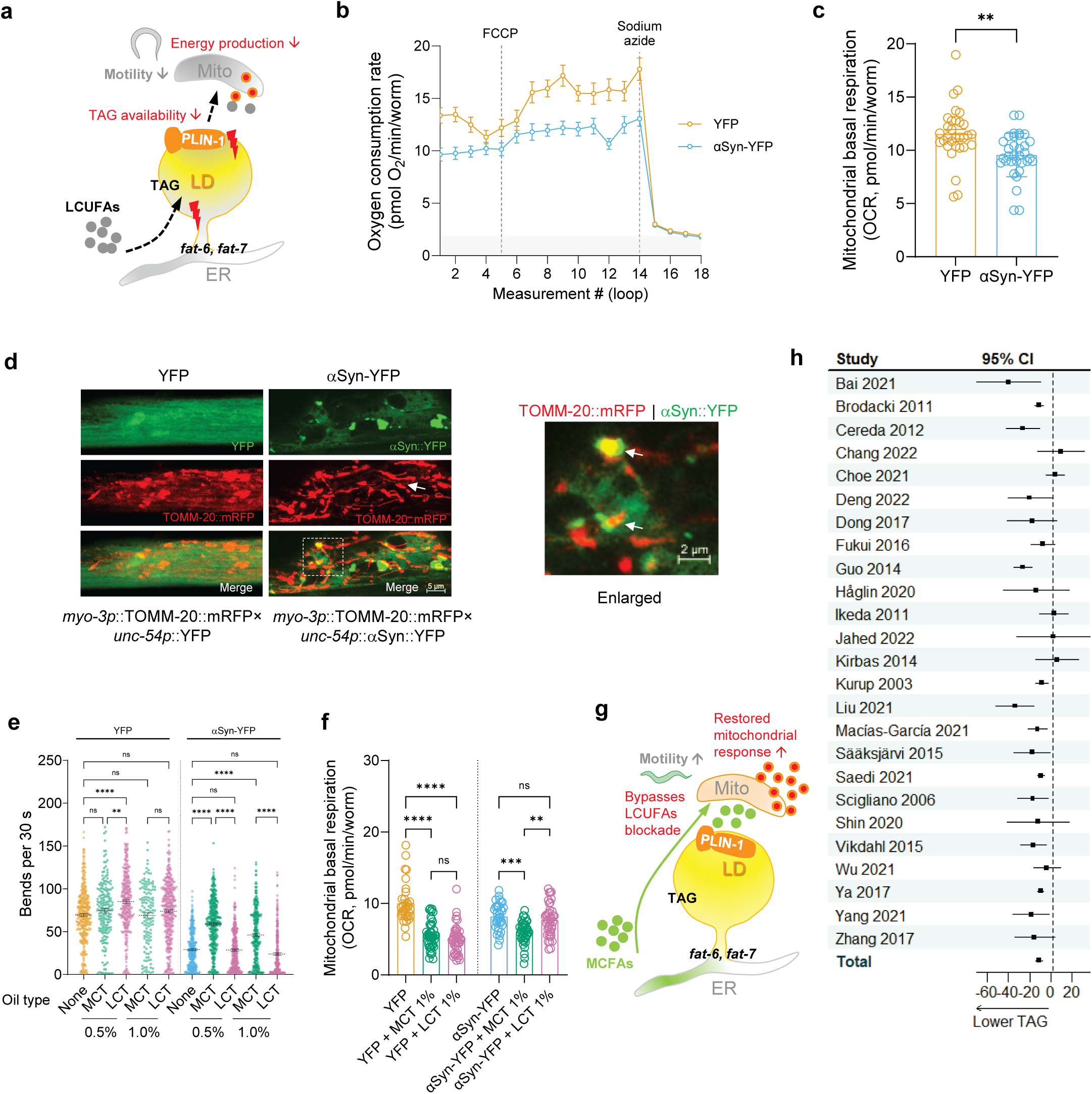
MCFAs supplementation bypasses LCUFAs synthesis to restore mitochondrial function and reduce αSyn toxicity. ***(a)*** Proposed mechanism illustrating *fat-6* and *fat-7* incorporate LCUFAs into TAGs for LD formation. In αSyn-YFP worms, disrupted LD biogenesis PLIN-1 mis-localization, impairs mitochondrial respiration by reducing TAG availability. ***(b)*** Oxygen consumption rate (OCR) measurement shows reduced basal mitochondrial respiration in day 5 of αSyn-YFP worms versus YFP controls (n=12 wells). ***(c)*** αSyn-YFP worms exhibit significantly diminished mitochondrial basal respiratory capacity compared to controls (** P<0.01, unpaired *t*-test). ***(d)*** Mitochondrial morphology assessment using TOMM-20::mRFP reveals increased fragmentation in αSyn-YFP worms (arrow) compared to YFP controls at day 5 of adulthood. ***(e)*** Dietary intervention with MCT but not LCT (olive oil), rescues motility defects of αSyn-YFP at day 5 adulthood (n>50 worms combined from 3 replicates; *P<0.05, ***P<0.001, one-way ANOVA). ***(f)*** The lowered basal mitochondrial respiration in αSyn-YFP worms when feeding 1% MCT indicates a reduced energy demand and a balanced energy supply in day 5 of αSyn-YFP worms compare to LCT feeding (** P<0.01, *** P<0.001, **** P<0.0001). ***(g)*** A proposed model shows that MCFAs supplementation via MCT feeding bypasses the impaired *fat-6* and *fat-7* mediated LCUFAs synthesis pathway and provides efficient substrates for a balanced mitochondrial respiration and energy production, resulting in rescued motility. ***(h)*** Forest plot of pooled results demonstrating significantly lower serum TAG levels in clinical PD patients (n=2,958) compared to healthy controls (n=10,112). All experiments were performed in triplicate. In ***(c)*** and ***(d)***, data were combined from three replicates.

As our data implicate LCUFAs in αSyn toxicity, we next asked whether bypassing LCUFA synthesis by dietary supplementation of medium-chain triglycerides (MCTs) could alleviate these defects. Unlike long-chain triglycerides (LCTs), MCTs, composed of C8–C12 MCFAs, are rapidly oxidized in mitochondria independently of the carnitine shuttle^33^. Indeed, motility in αSyn-YFP worms was significantly improved when fed 0.5% or 1.0% MCT compared to YFP controls, whereas feeding LCT did not improve motility (Fig 6c). In addition, mitochondrial respiration measurements revealed that αSyn-YFP worms respond to MCT feeding but not LCT, while mitochondria of YFP worms respond to both MCT and LCT feeding (Fig. 6e). When feeding 1% MCT, different from LCT, αSyn-YFP worms showed a lowered basal mitochondrial respiration. Although counterintui.ve, given that MCT and LCT feeding both lower respira.on of YFP animals, the lowering could reflect a metabolic shiA of the mitochondria, requiring less oxygen for using faEy acids as an energy source. In contrast to YFP worms, only MCT can be used in αSyn-YFP worms due to underlying lipid metabolic defects (Fig. 6f and Fig. S6c). In all, these results demonstrate that dietary MCFAs can bypass the LCUFA metabolic blockade, restore mitochondrial response, and rescue toxicity in αSyn-expressing worms (Fig. 6g).

Finally, we conducted a systematic analysis of 25 clinical studies involving 2,958 PD patients and 10112 healthy controls to evaluate potential relevance of our findings in humans (Fig. S7a). Indeed, we found that serum TAG levels were significantly lower in PD patients compared to healthy controls (Fig. 6h and Fig. S7b). Supporting our conclusion that reduced food intake does not lower TAG levels, a previous study also reported a reduced whole-body fat mass index in PD patients despite comparable dietary energy intake^34^. In all, decreased TAG levels appear to represent a conserved feature shared between *C. elegans* PD models and clinical PD.

## Discussion

αSyn pathophysiology is closely associated with disrupted lipid homeostasis but underlying cellular mechanisms are poorly understood^35^. Here, we identified a previously unrecognized conservation of TAG depletion between *C. elegans* models and PD patients. Intriguingly, systematic analysis of a series of clinical studies revealed consistently lower fasting serum TAG concentrations in PD patients compared to healthy controls. As αSyn condensates impair mitochondrial function, they promote toxicity by depleting TAGs and elevating LCUFA levels. Supplementation with MCTs, balances the energy demand and rescues pathological effects by bypassing the need for TAG mobilization. As TAG mobilization from LDs is essential for supplying FFAs required for beta-oxidation as well as for mitochondrial biogenesis^36^, our findings suggest that MCTs restore mitochondrial respiration response, thereby improving motility in our αSyn worms.

Among the TAG metabolic genes screened, suppression of LCUFAs biosynthesis significantly reduced αSyn toxicity. This protective effect aligns with previous studies attributing the benefit to the balance between saturated and unsaturated fatty acyl chains^6^. Stearoyl-CoA desaturase (SCD), the mammalian homolog of *fat-6* and *fat-7*, has been extensively studied in PD and related synucleinopathies^37^. Knockdown of *fat-6* or *fat-7* likely confers protection by limiting SCD-mediated desaturation of saturated fatty acids into their monounsaturated forms (C16:0 into C16:1 and C18:0 into C18:1). Although SCD inhibition-mediated low desaturation may decrease αSyn toxicity, we observed increased proportions of unsaturated PE in αSyn worms, suggesting that *fat-6* and *fat-7* knockdown may exert protective effects through multiple SCD-dependent mechanisms. These findings are consistent with the role of SCD in regulating the rations of phosphatidylcholine (PC) to PE^38^. In line with previous findings that *fat-6*-dependent products are essential for development^39^, *fat-6* knockdown reduced body size and modestly improved motility in control worms, while significantly lowering αSyn expression and TAG levels. These findings suggest that the protective effect of *fat-6* knockdown reflect systemic metabolic changes rather than αSyn-specific toxicity suppression. In contrast, *fat-7* overexpression significantly increased αSyn levels and severely impaired motility without affecting body size, consistent with a distinct and possibly more beneficial role for *fat-7* in modulating αSyn toxicity. Together, these results reveal an unrecognized functional divergence between the two SCD isoforms in αSyn toxicity. Further studies are needed to determine whether *fat-6* and *fat-7* act in a tissue-specific manner or under distinct physiological conditions. For example, *fat-7*, but not *fat-6*, is required for maintaining membrane fluidity under stress^40^. Additionally, whether the observed TAG depletion results from decreased *de novo* fatty acid synthesis or enhanced mobilization remains unclear.

We also observed pronounced mislocalization of PLIN-1 in αSyn-expressing worms. This phenotype may reflect a reduction in LD surface area due to TAG depletion, limiting PLIN-1 anchoring. As PLIN-1 facilitates FFA release from LDs for mitochondrial beta-oxidation, its mislocalization could impair TAG storage and lead to premature lipolysis. Notably, the neuronal αSyn(A53T) mutant, which induces more severe toxicity, caused a stronger reduction of TAG levels, further supporting the association between αSyn toxicity and disrupted TAG homeostasis. This finding is consistent with recent studies indicating that TAGs act as a critical energy reserve for neuronal synaptic function^41^, with efficient FFA mobilization from LDs beingessential for local mitochondrial energy production. Collectively, these findings suggest that αSyn condensation leads to a broader disruption of TAG homeostasis that extends beyond its immediate site of expression.

Interestingly, while αSyn inclusions occasionally colocalized with LDs at early developmental stages, we did not observe consistent colocalization at later stages, implicating mechanisms beyond direct LDs binding. For example, αSyn could bind FFAs or interact with TAG-metabolizing enzymes localized outside LDs^42^, thereby impairing LDs formation or turnover. In previous studies, αSyn was reported to inhibit TAG lipolysis by associating with LDs, leading to their accumulation^43, 44^. Consistently, we found that deletion of αSyn’s membrane-binding domain rescued defects in motility. In addition, we observed a co-localization pattern between αSyn and mitochondria suggesting that αSyn may impair mitochondrial function by interfering with energy supply. In all, these findings underscore the critical role of interactions with lipid-rich organelles in αSyn-mediated toxicity.

Our data also shows that αSyn condensates induce mitochondrial fragmentation, warranting further investigations of mitochondrial dynamics. Additionally, αSyn may exert localized effects at other organelle contact sites, such as mitochondria-associated ER membranes (MAMs)^45^. αSyn has been reported to disrupt ER-Golgi trafficking and to interact with both mitochondria and LDs^46, 47^. These interactions interfere with the ER-mediated conversion of DAG to TAG, resulting in excess FFA with potential toxicity. Sequestering excess FFAs into LDs reduces αSyn aggregation, while limiting the incorporation of unsaturated FFAs into TAG alleviates αSyn-induced toxicity^43, 48-50^. Collectively, these findings underscore the central role of LD imbalance in αSyn pathology, particularly involving alterations in fatty acyl chain length and saturation. Our study identifies TAG imbalance as a critical pathological feature, linking upstream αSyn-induced lipid alterations to downstream impairments in mitochondrial energy metabolism.

In the future, it will be important to elucidate the subcellullar alterations of specific TAG species and the underlying mechanisms involving organelle interactions, as well as the structural changes induced by elevated αSyn condensation. TAGs are hydrolyzed to release FFAs, which serve as essential substrates for energy production but also function as signaling molecules during tissue repair. It is also interesting to consider whether MCTs and other structurally related TAG species support mitochondrial ATP production. Such metabolic flexibility could offer a targetable therapeutic strategy for PD and related synucleinopathies, especially under conditions in which access to other energy sources is limited^41^. Overall, manipulating mitochondrial energetics represents a promising therapeutic approach for the treatment of PD and related disorders.

## Materials and methods

### C. elegans strains

Standard conditions were used for maintaining C. elegans strains at 20°C. Animals were synchronized by hypochlorite bleaching, hatched overnight in M9 buffer at 20 °C and subsequently cultured on NGM agar plates seeded with OP50. Worms were grown from L1 until day 5 of adulthood at 20°C. For ageing experiments, worms were transferred at L4 stage to NGM plates containing 5-Fluoro-2’deoxy-uridine (FUDR) to inhibit growth of offspring. All the strains used in this study are listed in Table S1.

### Oil Red O staining

To measure fat accumulation, synchronized adult animals were stained with Oil Red O (ORO)^51^. Briefly, 100 worms were washed three times with M9 and allowed to settle at the bottom by gravity. The supernatant was removed, and worms were fixed with 1% paraformaldehyde (PFA), dissolved in M9 at 1:1 ratio. The samples were left to rotate for 30 min at room temperature. They were then freeze-thawed three times and washed again with M9 three times to remove PFA. The worms were then resuspended in 60% isopropanol and incubated for 15 min at room temperature. After the worms were allowed to settle at the bottom, the isopropanol was removed, 1 mL of 60% ORO working solvent was added and the worms were incubated for 4 hours. The ORO was removed by washing three times with PBS. Worms were mounted on slides and imaged using stereomicroscope (Stereo DiscoveryV12; Carl Zeiss) with Axiocam 503 color camera.

### Food intake measurement

OP50::mCherry bacteria were cultured overnight in LB broth. A 0.2 mL aliquot of the harvested bacteria was spread onto a 6 cm agar plate and allowed to grow overnight. Worms were subsequently transferred onto the bacterial lawn. Five minutes after transfer, worms were picked and mounted on slides for observation under an imaging microscope. Images were captured using an AxioCam HRm camera and ZEN lite software. Fluorescence intensity was quantified and analyzed using ImageJ.

### Sample preparation for lipidomics

To analyse lipid composition using mass spectrometry, YFP (OW450) and αSyn-YFP (OW40) worms were collected at day 11 of adulthood. Synchronized animals were collected in M9 buffer in 15 mL tubes and washed at least four times, by allowing the worms to settle by gravity between washes to remove bacteria. Finally, worms were transferred to 1.5 mL tubes, frozen with liquid nitrogen and stored at -80°C until analysis.

### Lipidomics analysis

According to a previous study^52^, around ∼2500 worms were collected in a 2 mL tube. A group of mixed internal standards were used for correcting the variation in lipid extraction and analysis. A 5 mm steel bead and 1:1 (v/v) methanol:chloroform was added to each sample to 1.5 mL. Samples were homogenized using a TissueLyser II (Qiagen) for 5 min at 30 Hz and centrifuged for 10 min at 20,000 g. The protein concentration of the lysate was determined using a Pierce™ BCA protein assay kit (Thermo Scientific). The supernatant was transferred to a 2 mL tube and evaporated at 45°C. The dried lipids were redissolved in 50 μL chloroform/MeOH/H2O (60:30:4.5, v/v/v) and further diluted with 150 μL isopropyl alcohol (IPA):acetonitrile (ACN):H_2_O (2:1:1, v/v/v). Lipids were separated by using an Acquity UPLC CSH column (1.7 μm, 100 × 2.1 mm) on an Acquity UPLC system (Waters, Manchester, UK). Mobile phases consisted of 10 mM ammonium formate in water (eluent A) and 10 mM ammonium formate in methanol (eluent B). Linear gradient elution was as follows: 0–5 min from 50 to 30% eluent A, 5–15 min from 30 to 10% eluent A, and 15–25 min from 10 to 0% eluent A. This was followed by isocratic elution at 0% eluent A over the next 15 min. A conditioning cycle of 5 min with the initial proportions of eluents A and B was performed prior to the next analysis. The column temperature was set at 80 °C, and the flow rate was 0.5 mL/min. 2 μL of sample was injected randomly in positive and negative modes, respectively. Mass spectrometry detection was performed using a Synapt G2-Si high-resolution QTOF mass spectrometer (Waters, Manchester, UK). Nitrogen and argon were used as desolvation and collision gas, respectively. Data were acquired over the m/z range from 50 to 2000 Da in continuum and enhanced resolution modes, at an acquisition rate of 1 spectrum/0.3 s. The source temperature was set at 120 °C, the desolvation temperature at 600 °C, the cone voltage at 30 V, and the capillary voltage was 0.50 kV for ESI(+) and 0.70 kV for ESI(−). MS experiments were performed with MS^E^ mode. The LockSpray internal reference used for these experiments to allow operation of the instrument at high mass accuracy.

To reduce and analytical variation, each batch included randomization of the sequence order and measuring a quanlity control (QC) sample at regular intervals (every 6 samples) in both modes. The identified lipids were presented as sum of FA chains without precise structural identification. Data were filtered on the pooled QC samples that were included in each run (coefficient of variation < 30%). All detected metabolite intensities were normalized to appropriate internal standards, as well as total protein content in samples, determined using a Pierce™ BCA Protein Assay Kit.

### RNAi screening

RNAi screening experiments were performed on 6 cm NGM agar plates containing 1 mM isopropylthio-β-D-galactoside (IPTG) and 50 mg/mL ampicillin that were seeded with RNAi bacteria. RNAi plates were kept at room temperature for at least 3 days to allow the bacteria to produce dsRNA. First larval stage (L1) animals were grown on RNAi plates after hypochlorite bleaching and used for the experiments. At L4 stage after synchronization, add 1 mL FUDR (125 μg/mL) to the NGM plates to inhibit growth of offspring. The RNAi clones used were verified by sequencing the insert using L4440 primers and subsequent BLAST analysis to confirm the correct sequence. After a first round in which ∼62 genes were individually knocked down and motility was determined on day 5 of adulthood. The motility was analyzed by using wide field-of-view nematode tracking platform (WF-NTP) according to our previous study^28^.

### Lipid feeding and staining

The detergent Tergitol (type NP-40, Sigma-Aldrich) was added to a final concentration of 0.001% in NGM agar medium to facilitate lipid dissolution for both non-supplemented and supplemented plates after autoclaving. 6 cm NGM plates for Lipid feeding experiments were prepared as described, and OP50 and RNAi bacteria (for *fat-6* and *fat-7* RNAi) were used for all lipid supplementation experiments. A final concentration of 0.8 mM oleic acid (OA) or 0.5%-1% oil was used. L4440 bacteria expressing the empty vector (EV) or the appropriate RNAi were seeded onto plates containing the respective lipid at room temperature for 24–48 h before the addition of worms to ensure incorporation of lipids into feeding bacteria^39^.

### Bodipy staining

Muscle lipid droplet was visualized using Bodipy 558/568 C12 (4,4-Difluoro-5-(2-Thienyl)-4-Bora-3a,4a-Diaza-s-Indacene-3-Dodecanoic Acid, Molecular Probes) staining^53^. Bodipy 558/568 C12 was dissolved in DMSO to produce a 5 mM stock solution. The stock solution was then freshly diluted in M9 buffer (1: 1000) and spread on the surface of NGM agar plates seeded with OP50 to final concentration ∼125 nM and dried in the dark. The animals were then transferred to the Bodipy plates, incubated at 20 °C, and subjected to fluoresence microscopy.

### Western blot

Worms (approximately 20 per sample) were transferred directly into Laemmli buffer (5x) and boiled at 95°C for 10 minutes. The denatured samples were then loaded onto a 12% Tris-Glycine acrylamide gel and electrophoresed. Proteins were transferred onto a 0.2 mm nitrocellulose membrane. Membranes were blocked with 5% milk in PBS-T (0.1%) for 1 hour. For protein detection, primary antibodies against synuclein (1:1000 dilution) and GFP (1:1000 dilution) were applied in 5% milk, and anti-tubulin antibody (1:10000 dilution) in fresh 5% milk, and incubated overnight at 4°C. Antibody binding was visualized using Amersham ECL Prime Western Blotting Detection Reagent and imaged with the ImageQuant LAS400 Imaging unit (GE Healthcare).

### RNA isolation and quantitative PCR

To assess gene expression levels after RNAi knockdown, total RNA was extracted from *C. elegans* using TRIzol reagent according to the manufacturer’s instructions. RNA quality and concentration were determined using a NanoDrop 2000 spectrophotometer (Thermo Scientific). cDNA was synthesized from 1 µg of total RNA using the RevertAid H Minus First Strand cDNA Synthesis Kit with random hexamer primers. Quantitative real-time PCR (qRT-PCR) was performed using the Roche LightCycler 480 Instrument II (Roche Diagnostics) with 2 µL of 10-fold diluted cDNA as template and SYBR Green dye for detection. The PCR cycling conditions were as follows: initial denaturation at 95°C for 10 minutes; 40 cycles of 95°C for 15 seconds, 60°C for 30 seconds, and 72°C for 15 seconds; followed by a melting curve analysis with 95°C for 5 seconds, 65°C for 1 minute, and a gradual increase to 97°C. Relative transcript levels were calculated using a standard curve from pooled cDNA samples. Gene expression was normalized to the endogenous reference gene *pmp-3*.

### Microscopy analysis

Worms were anaesthetized using 50 mM levamisole and high magnification (40x objective) z stack images of the head region were obtained by using a Zeiss Axio imager 2 microscope or Confocal Microscopy (ZEISS LSM 900 with Airyscan). Lipid droplets were quantified manually, assisted by the Fiji analyze particle function applied to maximum intensity projections of the z stacks. At least 10 worms were quantified per time point per condition. Each experiment was performed in triplicate, unless stated otherwise.

### Mitochondria respiration meaurement

Oxygen consumption was assessed using a Bioscience Seahorse XF96 Flux Analyser (or XFe96) as previously described^32^. Synchronized 5-day-old worms were collected in M9 buffer and washed several times to remove bacteria and larvae. Each experiment was repeated three times.

### Meta-analysis of TAG levels in patients

This systematic review and meta-analysis were conducted in accordance with the Preferred Reporting Items for Systematic Reviews and Meta-Analyses (PRISMA) guidelines. To identify randomized controlled trials (RCTs) examining serum TAG levels in PD patients, we systematically searched PubMed, Embase, Web of Science, the Cochrane Library, and previous meta-analyses for relevant articles published up to November 25, 2022. The search strategy employed the following terms: [(((triacylglycerol and parkinson disease) OR (TAG and parkinson disease)) OR (TG and parkinson disease) AND (clinicaltrial[Filter] OR meta-analysis[Filter] OR randomizedcontrolledtrial[Filter] OR review[Filter] OR systematicreview[Filter])) OR (lipid and parkinson disease)]. Filters were applied to include only clinical trials, randomized controlled trials, and English-language publications, with no restrictions on population or publication date. Additionally, we manually screened reference lists from included studies and prior reviews to identify further relevant trials.

Following the retrieval process, studies were screened based on titles, abstracts, and full texts for eligibility. For dichotomous outcomes, we calculated pooled effect estimates using relative risk with 95% confidence intervals (CIs), applying the Mantel-Haenszel method, which is particularly suitable for small-scale studies. For continuous outcomes, we extracted either the mean change from baseline or post-intervention values along with standard deviations. Weighted mean differences were used when measurements were consistent, whereas standardized mean differences were employed for heterogeneous measures (e.g., differing scales or units). A random-effects model (DerSimonian–Laird method, based on the inverse-variance approach) was utilized to account for both within-study and between-study variability. Mean differences and SDs between baseline and endpoint were extracted for each trial. If these were not directly reported, the mean difference was calculated by subtracting baseline values from endpoint values and derived SDs from standard errors, 95% CIs, medians, or interquartile ranges where necessary. All statistical analyses were performed using Review Manager 5.4.

### Statistical analysis

All experiments were replicated three times, unless stated differently in the corresponding figure legends. Microscopy images were selected from random samples to avoid any artefacts and improve the robustness of the data. The investigators were not blinded to the experimental conditions during data collection and analysis. Statistical analyses were done in Graphpad Prism 9 (10.1.2). Based on the distribution of the data the appropriate test was selected (parametric or non-parametric). Principal component analysis (PCA) was performed using the FactoMineR package to visualize the relationships among transformed quantitative variables and to assess the distribution of the observed data. For group comparisons, an unpaired samples two-tailed t-test or one-way ANOVA was conducted, followed by Tukey’s multiple comparison test. Significant statistics (P values) is denoted as follows: ns, p > 0.056, *: p<0.05, **: p < 0.01, ***: p < 0.001, ****: p < 0.0001. Significance values are also indicated in the figures and figure legends. All data were visualized with Graphpad Prism 9 (10.1.2) or R (3.6.0).

## Acknowledgments

The *C. elegans* strains used in this work were provided by the Caenorhabditis Genetics Centre (University of Minnesota), which is funded by the NIH Office of Research Infrastructure Programs (P40 OD010440). The study was supported by the National Natural Science Foundation of China (Grant No. 32272302), and the grant from ParkinsonFonds.nl (to T.Z. and E.A.A.N., Grant No. 1907) and (to M.E.G. and E.A.A.N., Grant No. 1887), and the grant from MSD co-funded by PPP Allowance awarded by Health∼Holland, Top Sector Life Sciences & Health, to stimulate public-private partnerships (LKR201161 MSD ERIBA) and the European Union’s Horizon Europe call HORIZON-WIDERA-2023-ACCESS-02 under Grant Agreement No. 101159690 to A.H.B., and the European Union’s Horizon Europe research and innovation programme under the Marie Skłodowska-Curie Gant Agreement (No. 812830). N.T. and D.T. are supported by the Hellenic Foundation for Research and Innovation (H.F.R.I.) under the “1st Call for H.F.R.I. Research Projects to support Faculty members and Researchers and the procurement of high-cost research equipment” (Project Number: HFRI-FM17C3-0869, NeuroMitophagy), the European Commission Research Executive Agency Excellence Hub “CHAngeing” (GA-101087071) and the General Secretariat for Research and Innovation of the Greek Ministry of Development, the European Union – NextGenerationEU (Project Code: TAEDR-0535850, Acronym: BrainPrecision). We thank Theo S. Boer, Boshi Wang, and Pim de Blaauw for suggestions and support. We thank Christina Lilliehook for editorial assistance.

## Author contributions

T.Z., J.C.W., and A.H. performed the lipidomics and data analyses. T.Z., M.E.G., A.H., D.T., and L.G. performed the food intake experiments, the ORO experiments and the mitochondrial experiments. T.Z., R.I.S., and S.C. designed and performed the RNAi screening, and the genetic assays with the help of R.I.S., M.E.G., and S.C. and input from E.A.A.N., R.I.S., F.K., N.T., M.C., and M.R.H.F. Crosses of all the mutant strains and western blots were with the help of R.I.S., and M.E.G. and T.Z. performed the imaging analysis and feeding experiments, with the help and input of M.E.G., and A.H. analyzed all the data, created all the figures, and performed statistical analyses. T.Z., E.A.A.N., and M.E.G., and F.K. were extensively involved in discussing experiments and follow-up steps. T.Z. wrote the manuscript with the help of E.A.A.N., A.H., and M.E.G., and E.A.A.N., M.E.G., and A.H. revised the manuscript.

## Data and Materials availability

All materials generated in this study will be shared upon request.

## Raw data availability

All raw data are deposited in Figshare and can be accessed through the following link: https://figshare.com/s/3d98b330b47519b214f6.

---

**Supplementary Fig. 1.**
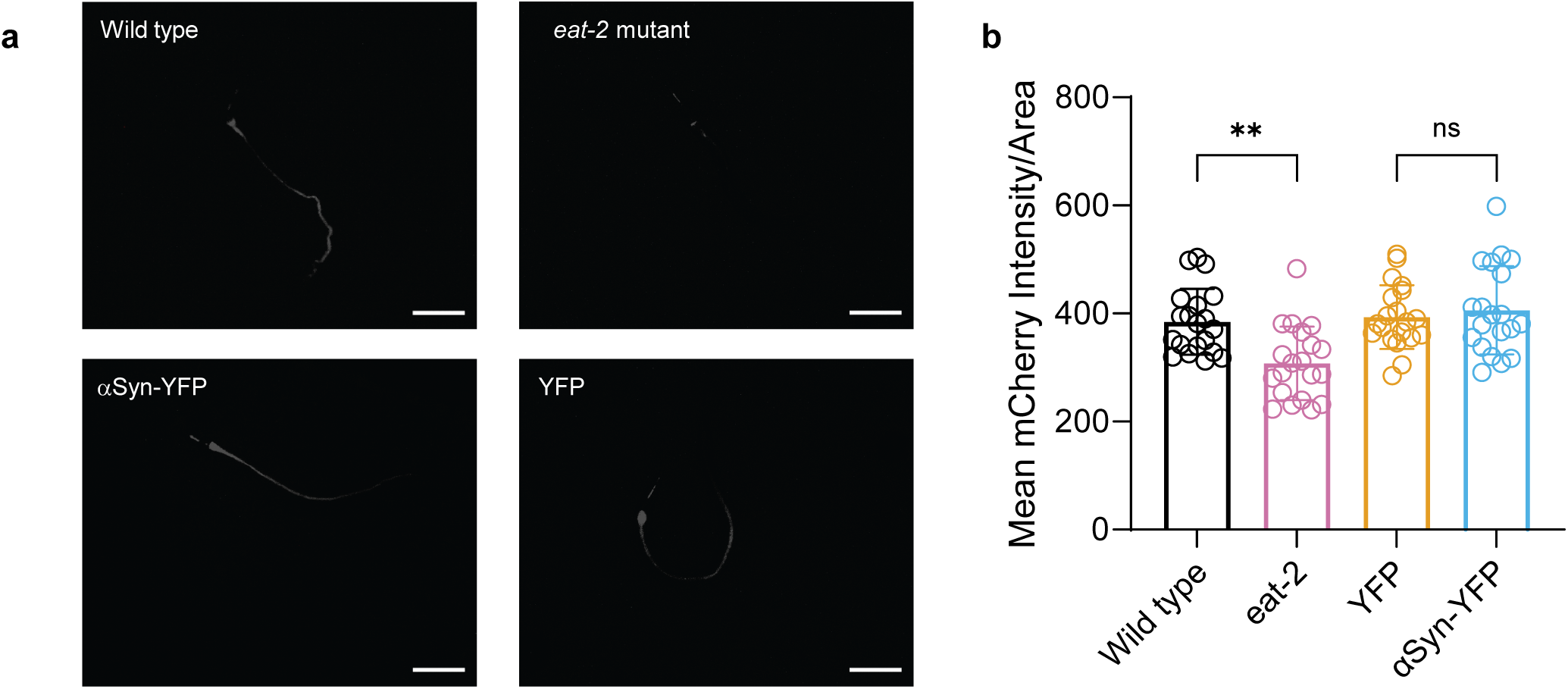
Comparable food intake between αSyn-YFP and YFP control worms. ***(a)*** Intestinal fluorescence in αSyn-YFP and YFP control worms fed mCherry-expressing *E. coli* OP50. ***(b)*** Quantification of OP50::mCherry fluorescence intensity in the intestinal lumen shows no significant difference in bacterial consumption between αSyn-YFP and YFP worms (Scale bar = 20 μm; n = 20 animals per group; **P < 0.01, one-way ANOVA). One representative image is shown. All experiments were performed in triplicate.

**Supplementary Fig. 2.**
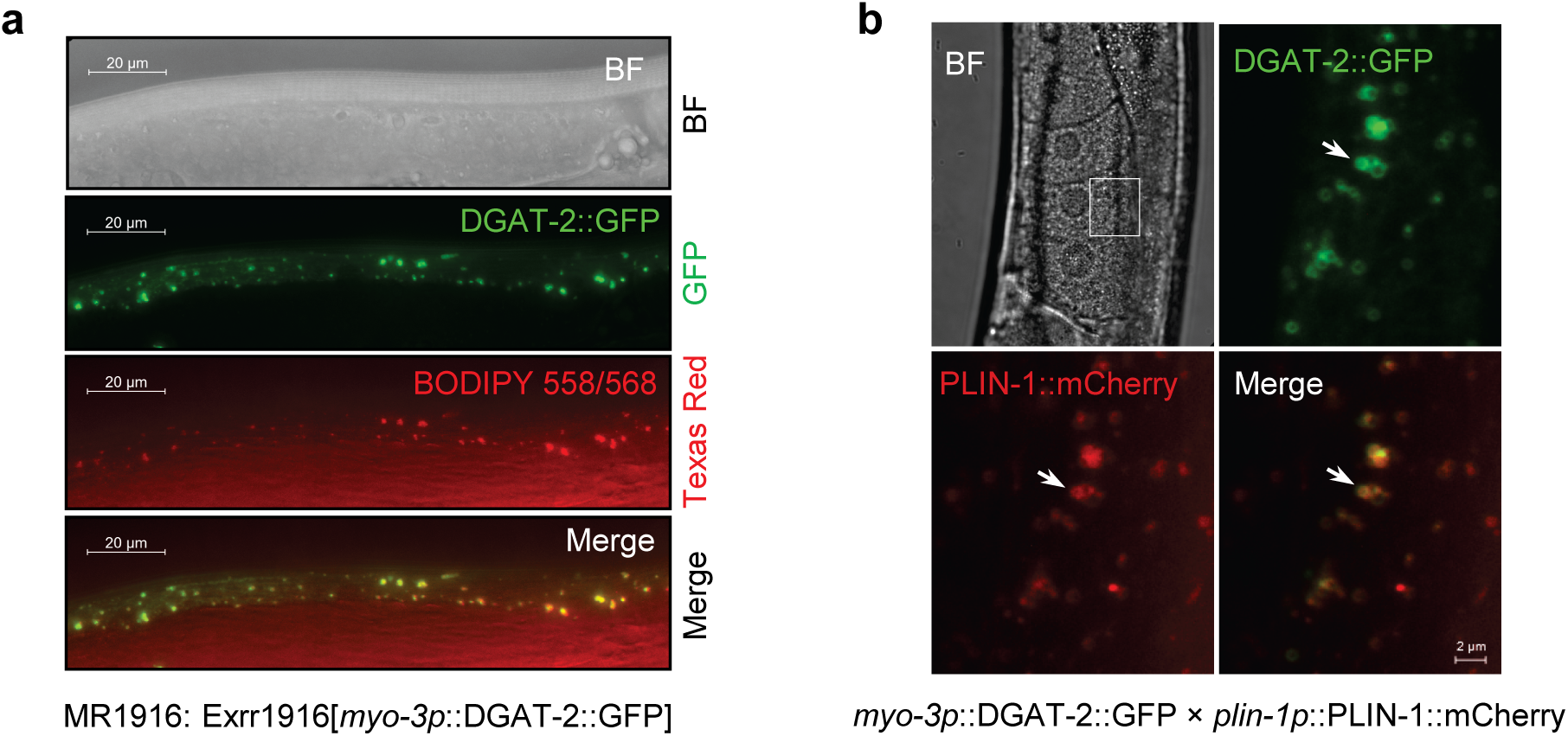
BODIPY-stained muscle LDs and PLIN-1::mCherry localized to muscle-specific LDs. ***(a)*** Muscle-specific LDs (stained with BODIPY 558/568 C12) colocalize with DGAT-2::GFP (expressed under the *myo-3* promoter in strain MR1916), confirming BODIPY-labeled LDs. ***(b)*** Muscle-specific co-expression of DGAT-2::GFP (green) and PLIN-1::mCherry (red) confirms PLIN-1 localization to muscle specific LDs. BF, bright field. All experiments were performed in triplicate.

**Supplementary Fig. 3.**
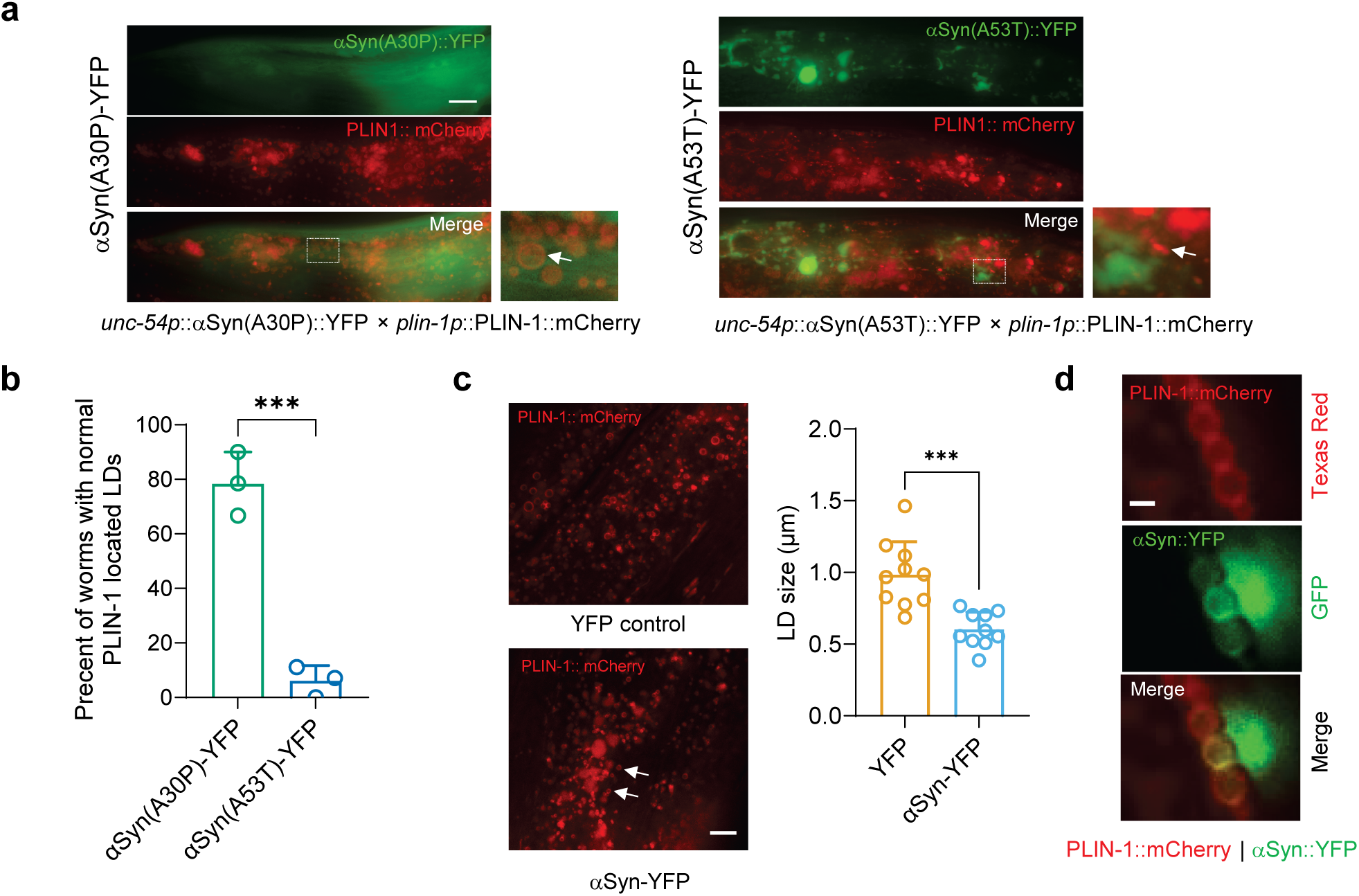
αSyn-YFP expression disrupts PLIN-1 localization and LD organizations. ***(a)*** αSyn(A53T)::YFP (green) compared to αSyn(A30P)::YFP induces abnormal PLIN-1 distributions and filamentous patterns (white arrow). Insets show magnified views of disrupted PLIN-1 (white-dotted boxes). Scale bars: 10 μm. ***(b)*** αSyn(A53T) significantly reduces the percentage of worms with normal PLIN-1-coated LDs at day 5 of adulthood (n = 15 animals per group; percentages derived from each biological replicate; ***P < 0.001, unpaired *t*-test. ***(c)*** αSyn-YFP worms exhibit significantly smaller LDs versus YFP controls at day 11 of adulthood (n=30 animals; ***P < 0.001, unpaired *t*-test). Scale bar: 5 μm. ***(d)*** Very little colocalization between αSyn::YFP with PLIN-1::mCherry was observed at younger stage (day 1 of adulthood). Scale bar: 1 µm.

**Supplementary Fig. 4.**
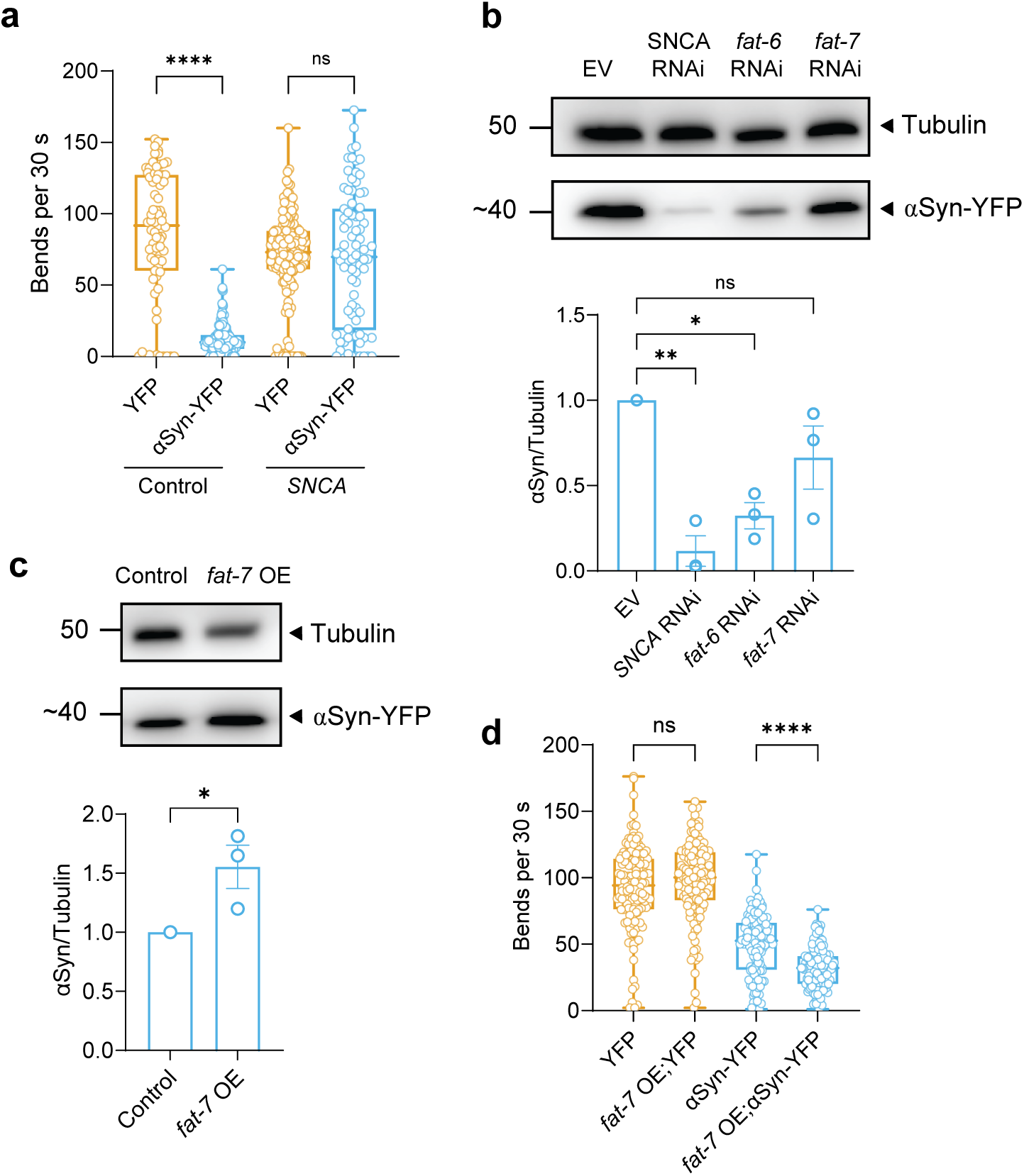
Differential effects of *fat-6* and *fat-7* modulation on αSyn toxicity. ***(a)*** Motility assay demonstrates that αSyn-YFP worms exhibit significantly reduced movement (body bends) compared to YFP controls at day 5 of adulthood, which is rescued by *SNCA* RNAi (n=50∼100 animals, ****P < 0.0001, one-way ANOVA). ***(b)*** Western blot analysis reveals *fat-6* RNAi significantly reduces αSyn protein levels in day 5 adult αSyn-YFP worms, while *fat-7* RNAi shows no significant effect (*P < 0.05, **P < 0.01, one-way ANOVA). ***(c) f****at-7* overexpression (OE) increases αSyn protein levels in αSyn-YFP worms (*P < 0.05, unpaired t test). ***(d)*** *fat-7* OE exacerbates motility defects in αSyn-YFP worms (n=50∼100 animals, ****P < 0.001, One-way ANOVA test). All experiments were performed in triplicate.

**Supplementary Fig. 5.**
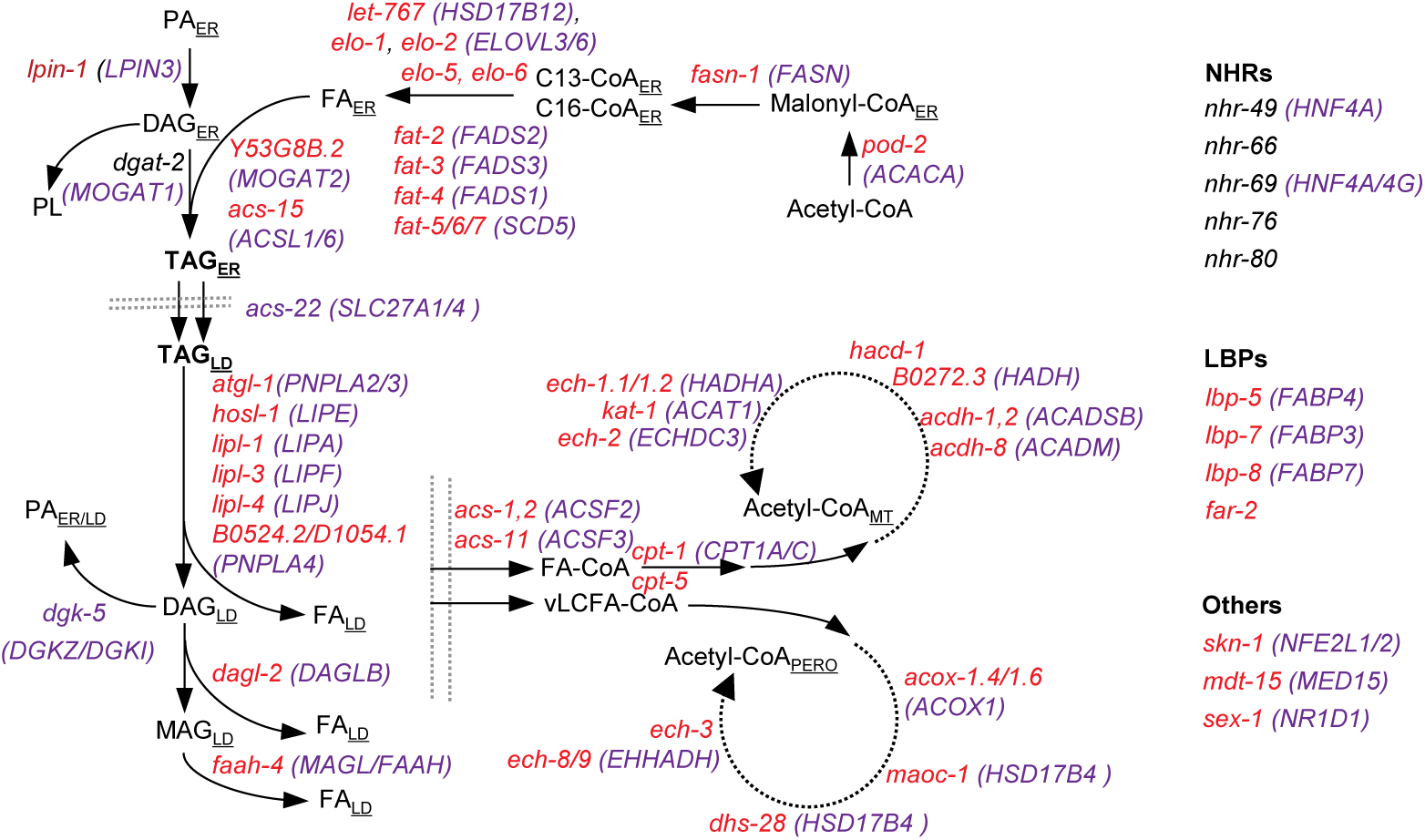
The lipid metabolism pathway in *C. elegans*. Human homologs of worm genes (red) are shown in purple.

**Supplementary Fig. 6.**
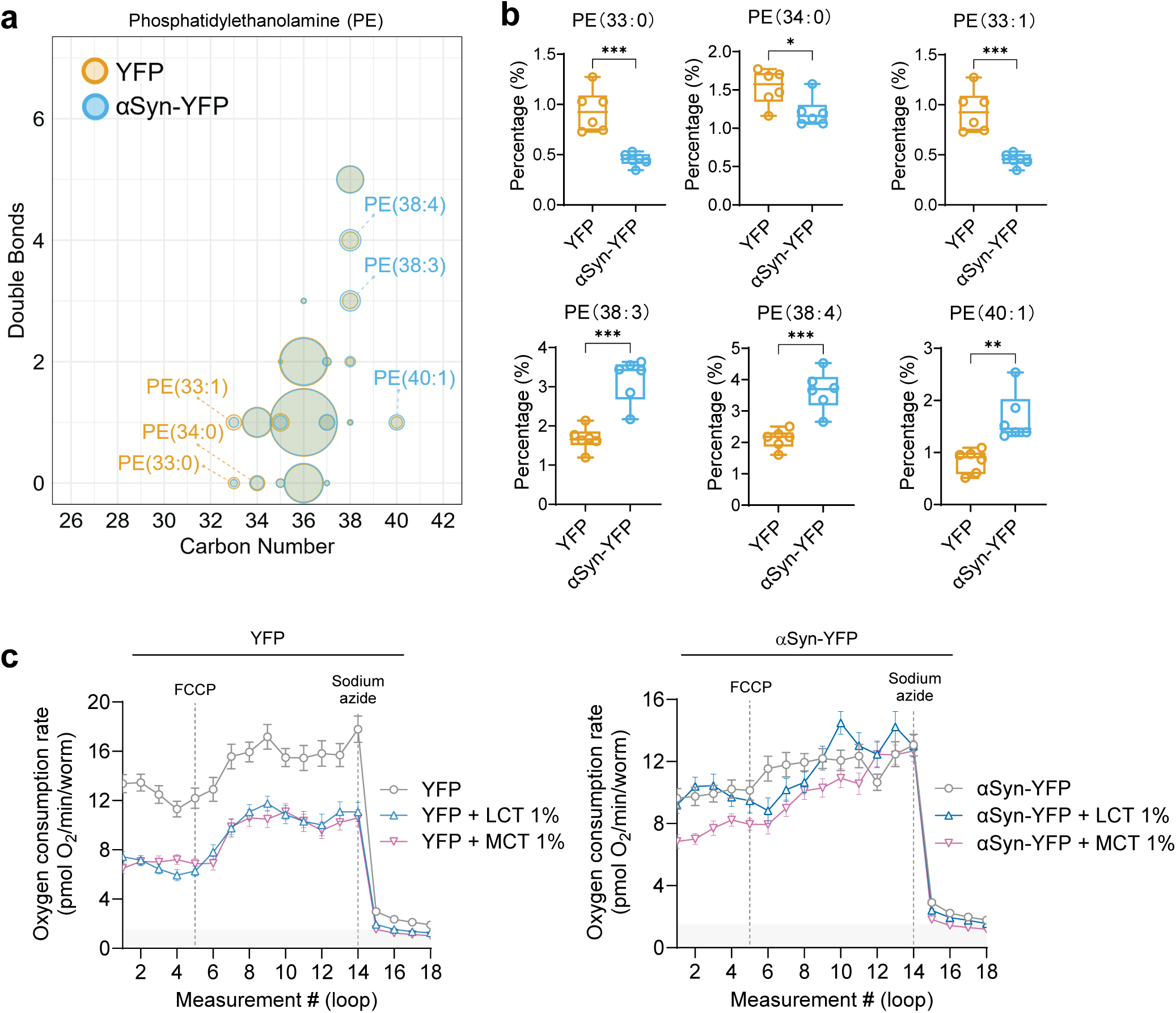
αSyn induces reservation of higher unsaturation of fatty acyl chains in phosphatidylethanolamine. ***(a)*** Lipidomics demonstrates increased unsaturation index (blue) in phosphatidylethanolamine (PE) species from αSyn-YFP versus YFP worms. ***(b)*** Percentage of six predominant PE molecular species shows higher unsaturated composition in αSyn-YFP worms compared to YFP controls (n=6). ***(c)*** Oxygen consumption rate (OCR) measurement shows αSyn-YFP worms specifically response to MCT feeding but not LCT feeding in day 5 of adulthoods, while YFP worms response to both MCT and LCT feeding (n=12 wells).

**Supplementary Fig. 7.**
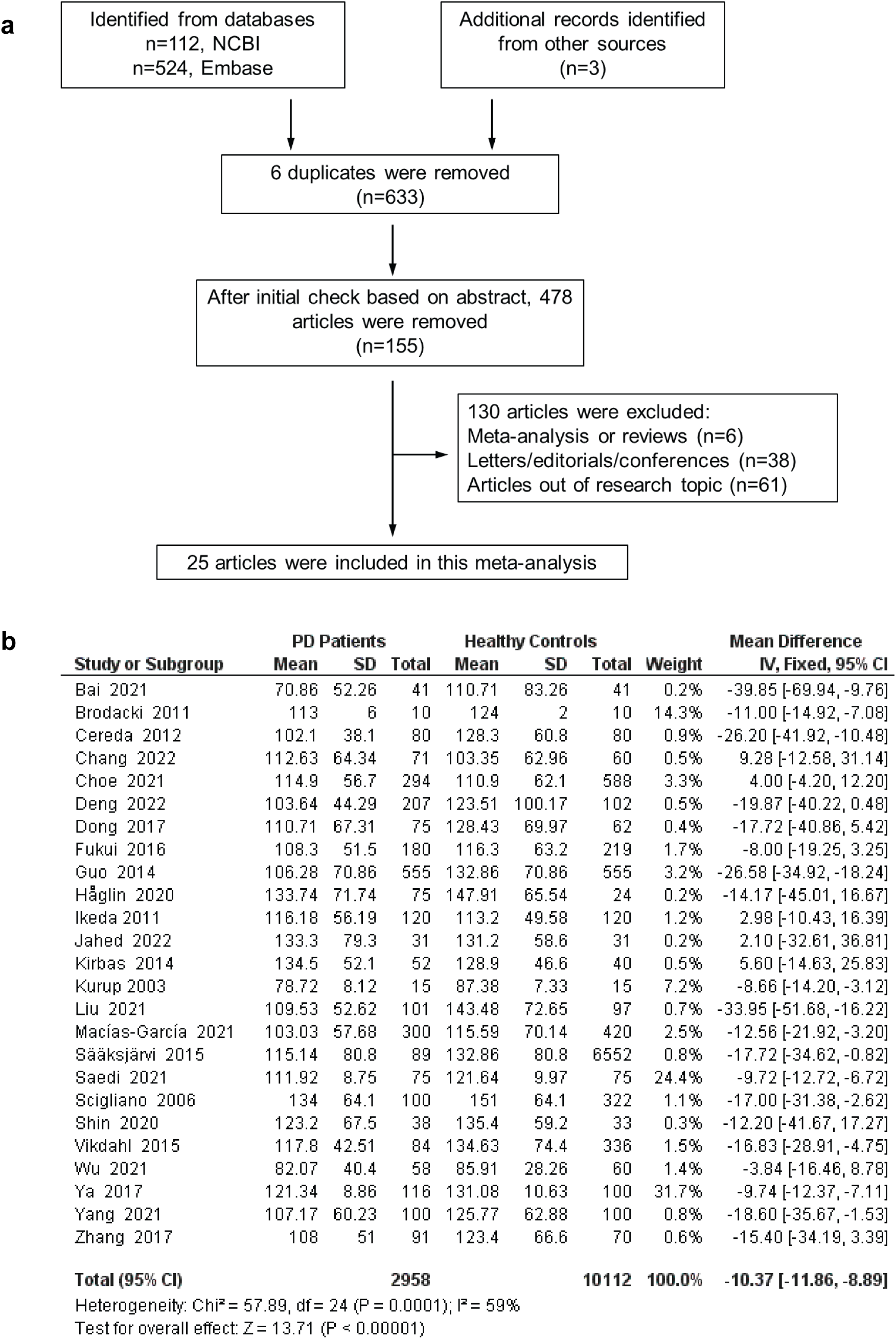
Reduced serum TAG levels in patients with Parkinson’s disease based on a systematic meta-analysis. ***(a)*** Flow diagram of study selection process for the meta-analysis, showing identification, screening, eligibility assessment, and final inclusion of studies. ***(b)*** Forest plot of pooled results demonstrating significantly lower serum TAG levels in PD patients (n=2,958) compared to healthy controls (n=10,112). The standardized mean difference (MD) with 95% confidence intervals is shown for each study and overall effect size (random-effects model).s

